# Protein-protein interaction prediction via structure-based deep learning

**DOI:** 10.1101/2023.05.27.542552

**Authors:** Yucong Liu, Zhenhai Li

## Abstract

Protein-protein interactions (PPIs) play an essential role in life activities. Many machine learning algorithms based on protein sequence information have been developed to predict PPIs. However, these models have difficulty dealing with various sequence lengths and suffer from low generalization and prediction accuracy. In this study, we proposed a novel end-to-end deep learning framework, RSPPI, combining Residual Neural Network (ResNet) and Spatial Pyramid Pooling (SPP), to predict PPIs based on the protein sequence physicochemistry properties and spatial structural information. In the RSPPI model, ResNet was employed to extract the structural and physicochemical information from the protein 3D structure and primary sequence; the SPP layer was used to transform feature maps to a single vector and avoid the fixed-length requirement. The RSPPI model possessed excellent cross-species performance and outperformed several state-of-the-art methods based either on protein sequence or gene ontology in most evaluation metrics. The RSPPI model provides a novel strategic direction to develop an AI PPI prediction algorithm.

## Introduction

Proteins serve as the basic building blocks playing specific roles in living cells, such as cell adhesion, signal transduction, post-translational modification, etc. [1, 2]. Generally, a biological process is accomplished by the cooperation of multiple proteins through transient protein-protein interactions (PPIs) [3]. High-throughput experimental methods have been widely used to determine PPIs, such as yeast two-hybrid screens [4], protein chips [5], and surface plasmon resonance [6]. However, experimental methods remain expensive, labor-intensive, and time-consuming. Therefore, the computational method has emerged in PPI studies. With the advance of machine learning and the accumulation of tremendous knowledge of protein and PPI information, large numbers of computational methods have been developed for predicting PPIs based on various data types, such as protein sequence [7] and protein secondary structure [8, 9].

The primary structures of the majority of proteins have been sequenced and stored in the UniProt database [10]. Thus, there is a longstanding interest in using sequence-based methods to model and predict protein interactions [11-13]. Several sequence-based methods have been developed to predict PPIs. For instance, Hashemifar et al. [14] proposed a deep learning model, DPPI, using a Convolutional Neural Network (CNN) and random projection modules to predict PPIs. DPPI achieved high prediction accuracy and could predict homodimeric interactions. Yang et al. [15] proposed a model based on a CNN architecture and protein evolutionary profiles. Their model showed superior performance and outperformed several other human-virus PPI prediction methods. However, the models based on the protein sequence showed low ability in generalizability [15, 16].

In addition, proteins have a higher chance to interact when localized in the same cellular component, or when sharing a common biological process or molecular function. Accordingly, several methods predict PPIs from the gene ontology (GO) annotations and semantic similarity of proteins [17]. Armean et al. [18] combined GO annotations with SVM for PPI prediction and other models employed in PPI prediction tasks including Bayesian classifiers [19] and random forest [20].

The PPI is mediated by non-covalent interactions, including electrostatic interaction, hydrophobic interaction, and hydrogen bonding, etc. In addition, the PPI is regulated by the fitness of the protein surfaces. Therefore, the structure of the protein and the physicochemical property of the surface amino acids together play an important role in PPI. However, the sequence-based and GO-based methods are incapable of incorporating protein spatial structure information. Thus, the structure-based model is needed for better learning PPI features. Currently, there are a few structure-based models for predicting PPIs. For example, Struct2Net [21] threads protein pair sequences to the protein complex in the Protein Data Bank (PDB) and searches for the potential match, and then predicts the PPI [22]. Cai et al. proposed a support vector machine (SVM) model to predict PPIs by analyzing the protein secondary structure [23]. Regrettably, these models based on protein structures required the corresponding homologous templates in PDB or ignored the global structure and physicochemical properties of proteins.

Herein, we designed a feature-extracting strategy, which converts 1D physicochemical properties and 3D structure information to 2D feature maps. The 2D feature maps were then fed to a newly proposed PPI deep learning framework, RSPPI. The RSPPI incorporated three ResNet blocks [24], an SPP layer [25], and a fully connected layer. The ResNet was used to extract the structure and physicochemical information; the SPP layer [25] was used to handle the problem of variable length of protein sequences; the fully connected layer was used to fuse the pairwise protein feature vectors and give the prediction of the interaction. Our model bridged sequence and structure information and transformed these features into 2D maps, which could be trained with ResNet. As RSPPI is based on structure and physicochemical features, the model does not rely on the homologous template of the PDB database. Moreover, thanks to the SPP layer our model does not need to preprocess the protein sequence into a fixed length, which may potentially introduce artifacts. These advantage of the model guarantees the outstanding performance and excellent cross-species predicting capability of the RSPPI model.

## Materials and methods

### Overview

We introduce an end-to-end deep learning framework, RSPPI, for identifying PPIs. The PPI prediction task is a binary classification problem. The amino acid physicochemical properties (including hydrophobicity and charge) and protein spatial structure (represented by distance map) were used as input in our RSPPI model. The model includes two parts: the data-preprocessing module and the prediction module (Fig. 1). In the data-preprocessing module, the 2D feature matrixes were generated from the physicochemical properties and spatial structure information of the protein. The 2D distance map, hydrophobic map, and charge map can be considered as multiple channels of an image with protein features. Inspired by the image recognition algorithms, we designed and trained a deep learning network. In the prediction module, the ResNet combined with an SPP layer was employed to learn the PPI features. Then, the pairwise protein representations are concatenated and fed into the designed fully connected layer to predict the interaction probability of the protein pairs.

**Figure 1.**
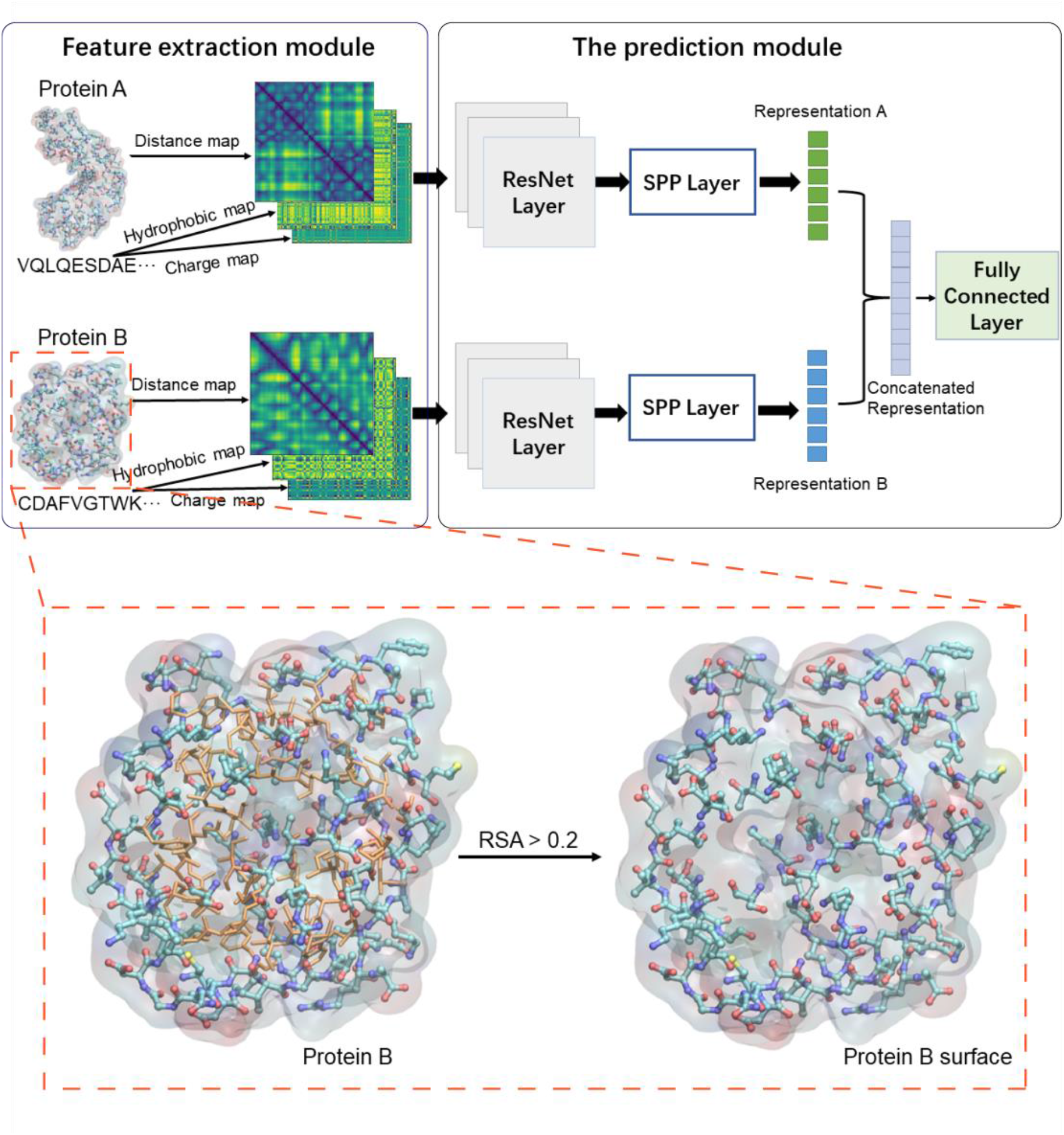
The overall architecture of the RSPPI model. All protein structures were pretreated by selecting the amino acids on the surface and then fed to the feature extraction module.

### Feature extraction module

The amino acids located on the surface of a protein have a prominent contribution to the protein interaction. Thus, the amino acids on the surface were carefully selected by calculating the relative solvent accessibility (RSA) [26]. The RSA of amino acids in the protein was calculated by the DSSP module of Biopython [27]. The amino acids with RSA exceeding 0.2 were defined as the surface amino acids [28]. A shell of the protein was then built on the selected surface amino acids and was used in the future data preprocessing (the inset in Fig. 1). A two-dimensional (2D) distance matrix of the protein shell was obtained by calculating the minimum heavy atom distance between residues, which represents the spatial structure information of the protein. As the amino acids in close contact have a greater chance to coordinate a specific interaction, we took more care of the amino acids which are close in space. Therefore, the protein distance map was further converted into an adjacency matrix. Any value less than or equal to a chosen cutoff of 14 Å was replaced by 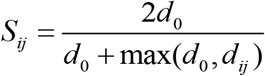, where *d*_*ij*_ is the heavy atom distance between the *i*th and *j*th residue, and cutoff distance *d*_*0*_ is set to 4 Å. Any value greater than the cutoff was set to 0 [29]. Therefore, the values of the adjacency matrix were in the range of 0 to 1.

The physicochemical properties of amino acids affect inter-protein interactions, such as their hydrophobicity and electrostatic charge. Hydrophobicity is a measure of how strongly the side chains of the amino acids in an aqueous environment are excluded by the water molecules. The hydrophobic amino acids of a protein tend to form hydrophobic interactions to reduce the interface with water or other polar molecules. The hydrophobicity is usually characterized by the hydrophobic moments (HM), ranging from -7.5 for Arg to 3.1 for Ile. The larger value HM indicates the amino acid possesses a higher hydrophobicity. As the hydrophobic interaction depends on the hydrophobicity of a pair of amino acids, the hydropathy compatibility index (HCI) was used to evaluate the hydrophobic interaction of amino acid pairs [30]. The HCIs of any two residues were calculated as:

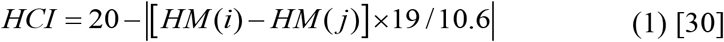

Where *HM* (*i*) and *HM* (*j*) are the HM of the residue *i* and *j*.

The electrostatic charges of amino acids define the electrostatic interactions. The residues with opposite charges attract each other, while residues with the same charge repel each other. A simple characterization of electrostatic properties is the isoelectric point (pI). The net charge of an amino acid varies with the pH of the environment owing to the gain or loss of protons. PI is the pH at which the net charge of the amino acid is 0. For instance, the negatively charged amino acids, Asp and Glu have PIs of 2.7 and 3.2, respectively, and the positively charged amino acids, His, Lys, and Arg, have PIs of 7.5, 9.7, and 10.7, respectively. The rest of the amino acids have PIs at ∼6, so they are considered to carry no net charge at the physiological condition. Similar to the HCL of the hydrophobicity, the charge compatibility index (CCI) was used to evaluate the electrostatic interaction of amino acid pairs [30]. The CCIs of any two residues were calculated as:

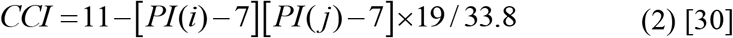

Where *PI* (*i*) and *PI* (*j*) are the PI of the residue *i* and *j*.

With the HCI and CCI, two more physicochemical feature maps of the protein surface shell could be constructed. Similar to the distance map, 2D hydrophobicity and electrostatic matrixes of the protein surface shell were built by assigning the corresponding element of the matrixes with the HCI and CCI value.

### Prediction module - ResNet layer design

Our network consisted of three ResNet layers with the same architecture (Fig. 3). In each layer, there were two ResNet blocks: a 1 × 1 Conv ResNet block and an identity ResNet block, respectively (Fig. 2). In both blocks, the feature maps firstly underwent convolution processing, then followed by an instance normalization [31], and eventually was activated via an exponential linear unit (ELU) activation layer [32]. These steps were repeated once again in the ResNet blocks. In the Convolution ResNet block, the size of the feature maps is halved owing to the 3 × 3 convolution kernel, while the number of output channels of the feature maps is doubled. Therefore, the original *x* had to be transformed to *x’* with an additional 1 × 1 Conv before adding up to the output *F’*(*x*). In contrast, thanks to the padding operation, the number of output channels and the size of the feature maps remained unchanged in the identity ResNet block. Therefore, the original *x* is added to the output *F*(*x*) at the end of the identity ResNet block. A kernel size of 3 × 3 was used in all the convolution operations. The filter number of the convolution operations in each ResNet layer was set to 16, 32, and 64, respectively. The ELU activation function was used as the activation layer because the ELU activation function is more effective than the standard ReLU in the ResNet [32]. Consequently, the feature maps of output changed from 3 × L × L to 64 × (L/8) × (L/8), where L is the protein sequence length.

**Figure 2.**
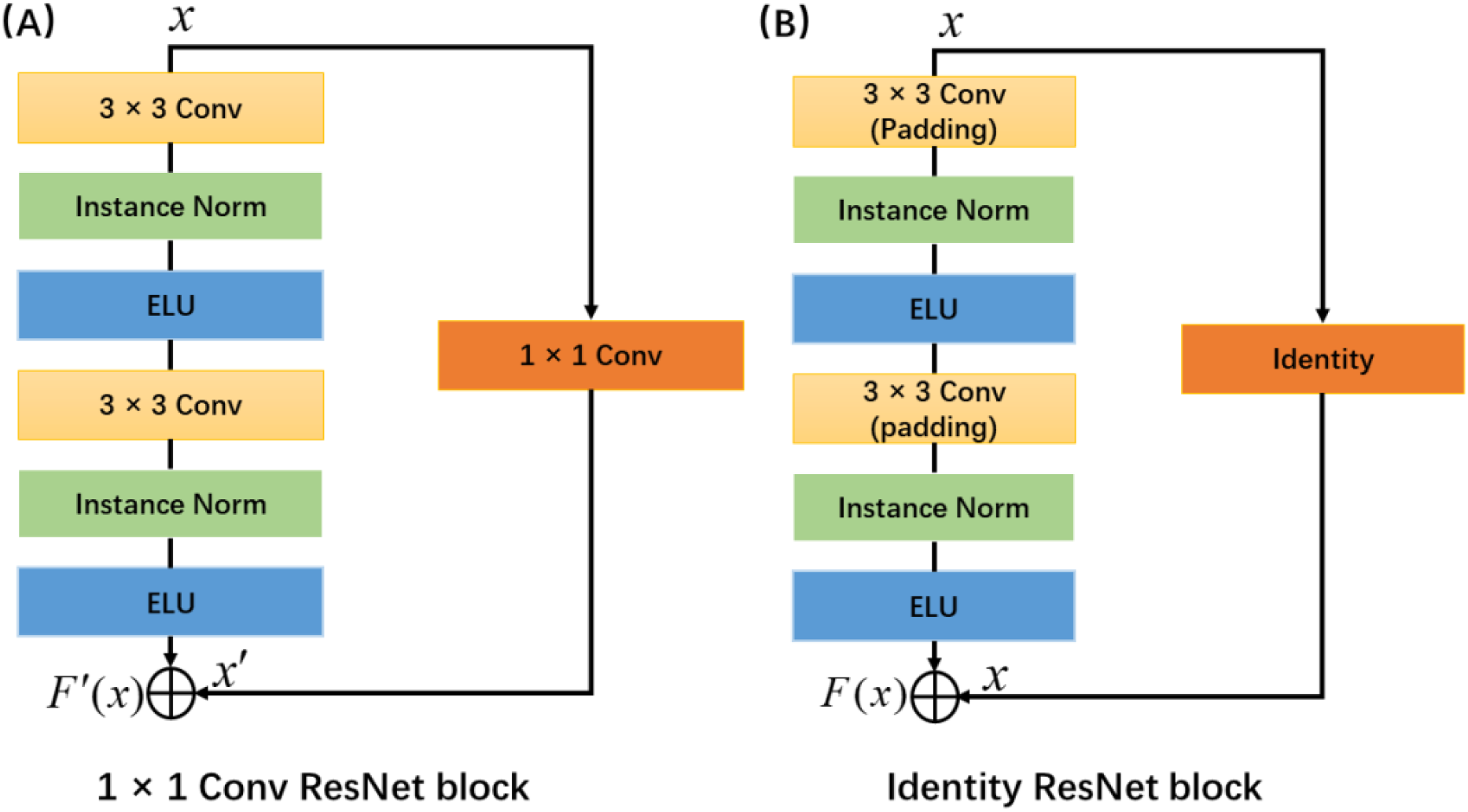
The framework of the convolution (A) and identity (B) ResNet blocks.

**Figure 3.**
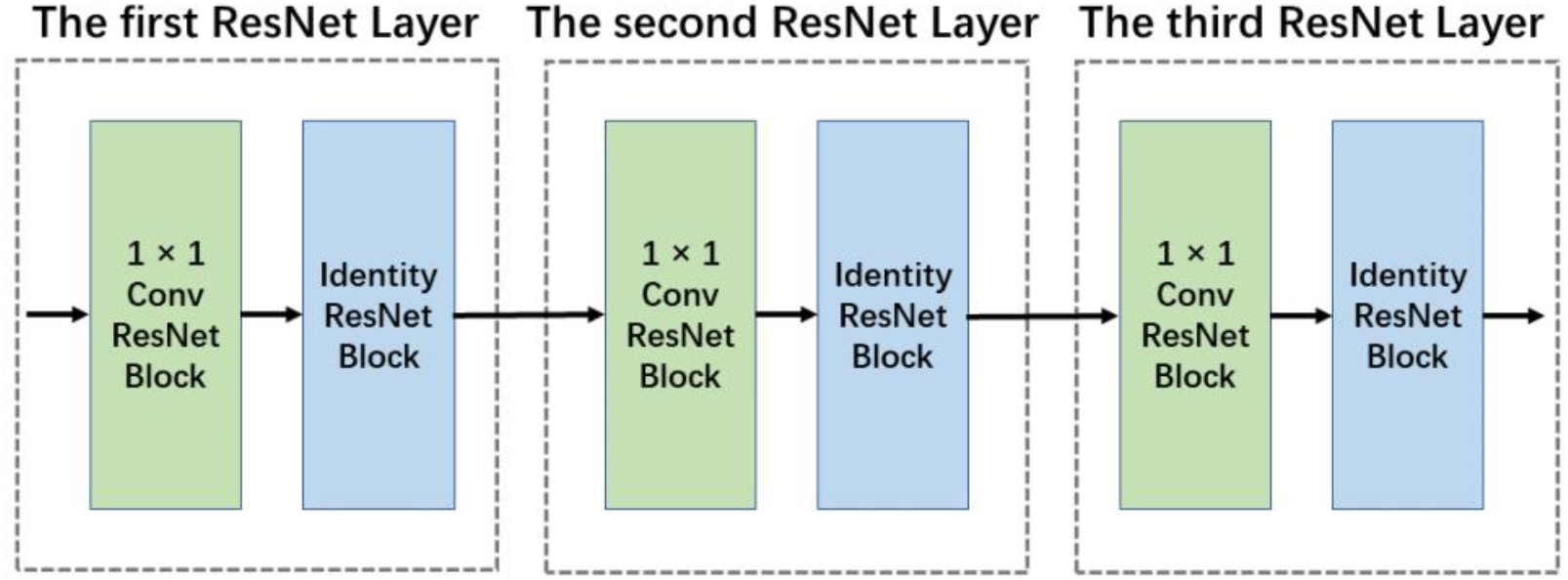
The framework of the ResNet Layer. Conv indicates the convolution kernel.

### Prediction module - Spatial pyramid pooling layer design

The fully connected layer requires a fixed-size input image. Therefore, a fixed length of the protein sequence is required to utilize the current CNNs with a fully connected layer for the PPIs prediction. Thus, truncating and zero-padding techniques for the lengthy proteins and short proteins, respectively, were widely used to unify the protein length. However, both truncating and zero-padding may introduce artifacts to the protein feature and influence the recognition. To bypass the fixed-length requirement, a four-layer SPP network was used after the ResNet layers. In the SPP network used here, four pooling operations were performed on the ResNet output feature maps. Four pooling grids of 4 × 4, 3 × 3, 2 × 2, and 1 × 1 were used for four pooling operations, respectively. As a result, four vectors with lengths of 16, 9, 4, and 1 were obtained from the four pooling operations, respectively. Considering the output of the ResNet is 64 feature maps, with the four pooling operations, 256 vectors were extracted (Fig. 4). By concatenating all the 256 vectors, a vector with a length of 1920 was obtained regardless of the size of the input protein.

**Figure 4.**
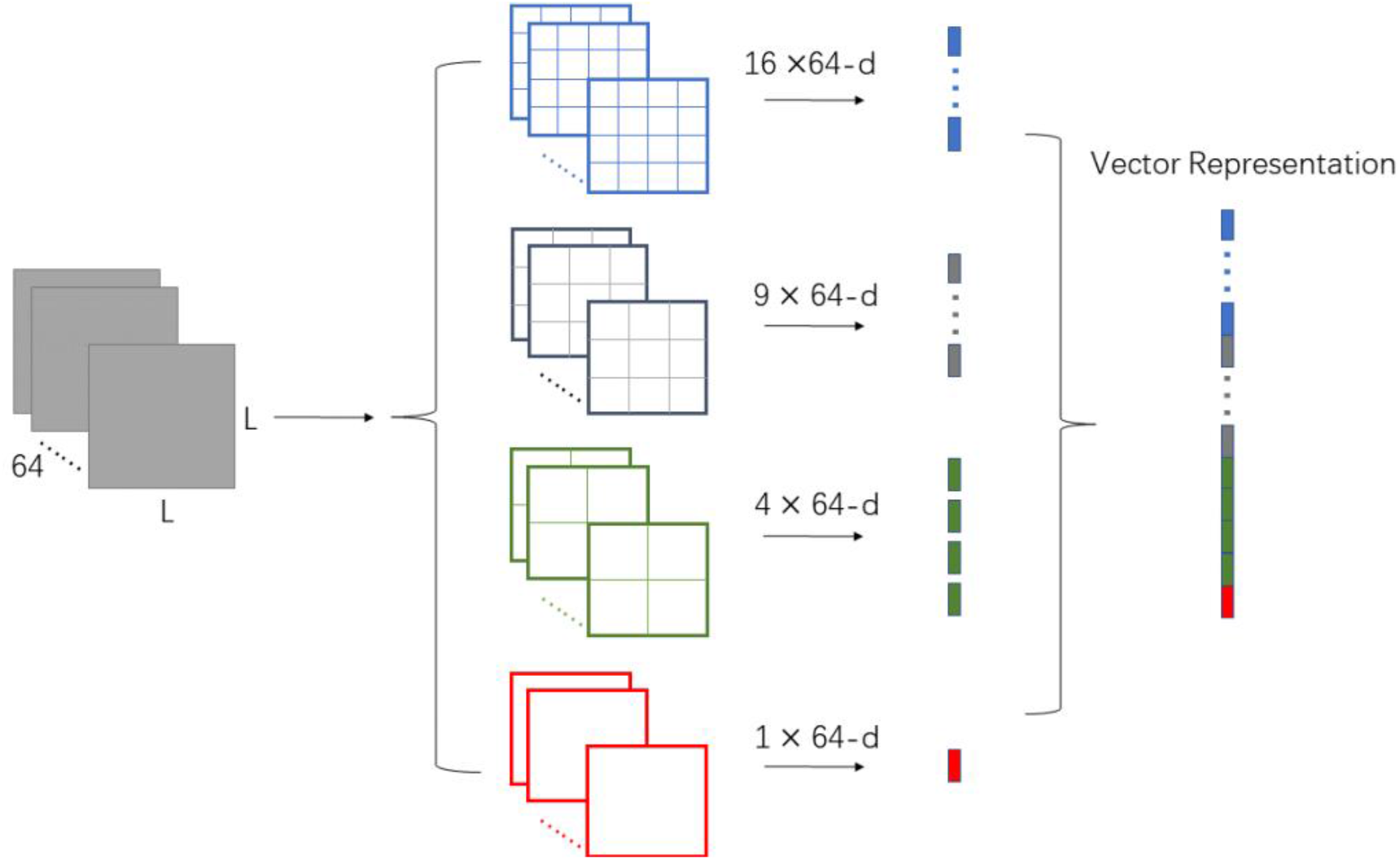
The architecture of the Spatial Pyramid Pooling Layer.

### Prediction module - Feature fusion layer design

To predict the interaction between two proteins, the pairwise protein vectors with the length of 1920 were extracted from the SPP network, respectively, namely *Z*_1_ and *Z*_2_.

We compute the difference and product of *Z*_1_ and *Z*_2_ to fuse the two protein features according to the following equations:

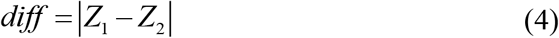

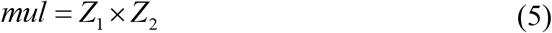

where *diff* indicates the element-wise difference and *mul* indicates the element-wise product. Then, the vector representations of *diff* and *mul* were concatenated to represent the pairwise protein feature, then the concatenated vector with 3840 dimensions fed into a fully connected layer. The fully connected layer stacks three linear layers followed by three ReLU activation functions, respectively, and an output layer followed by the Sigmoid activation function. The lengths of the linear layers and output layer were set to 3840, 960, 480, and 1 (Fig 5). BCELoss was used as a loss function to train the model, which was defined as:

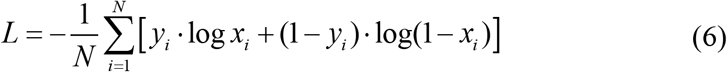

where *N* is the total number of protein–protein samples in the training dataset. *Y*_*i*_ and *x*_*i*_, respectively, were the true target and the predicted score of the *i*th sample.

**Figure 5.**
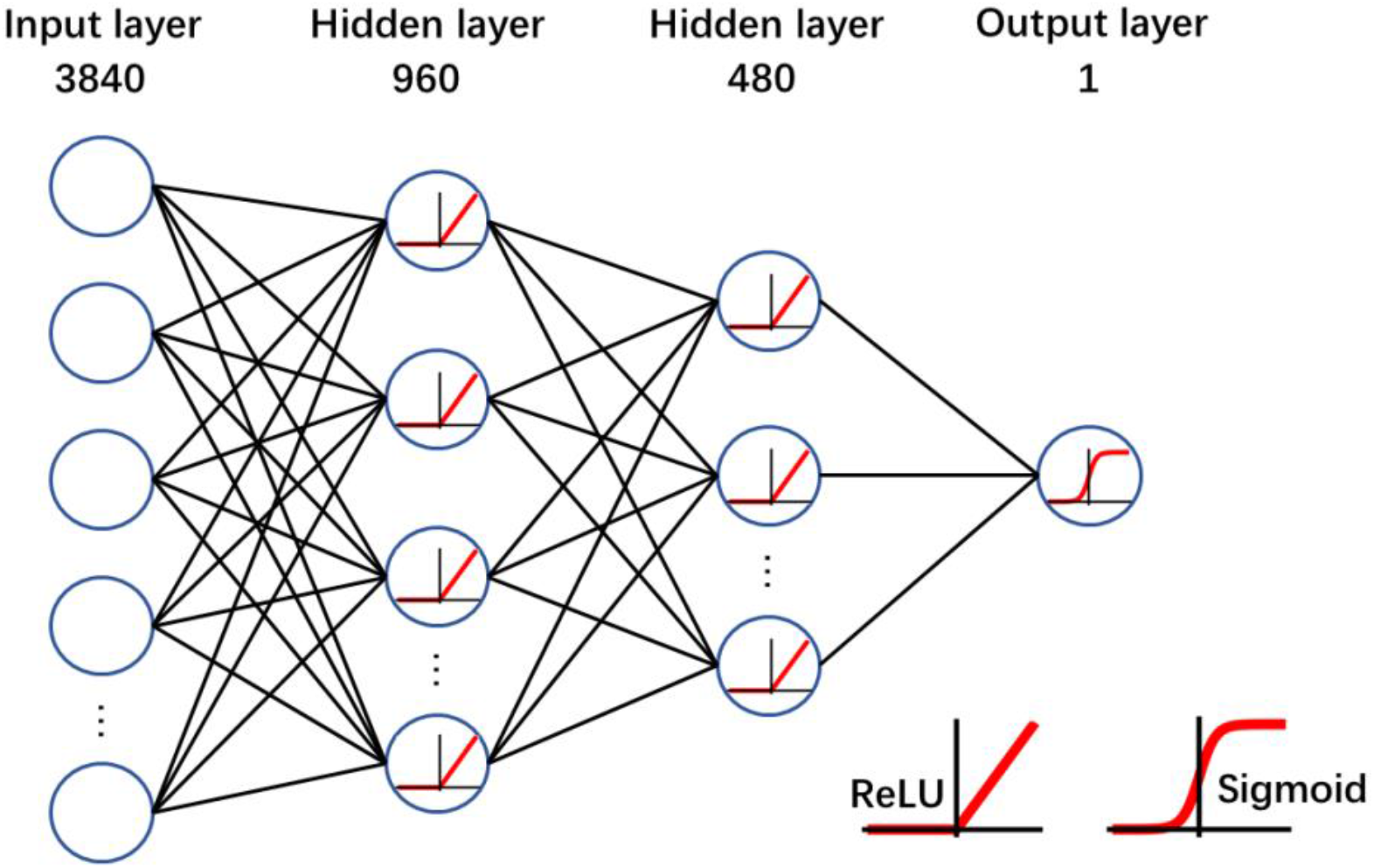
The architecture of the fully connected layer.

### Dataset preparation

As the PPI is the training purpose, the dimers stored in the Protein Data Bank (PDB) were collected to build the training dataset. The species with fewer structures were screened out. Thus, the dimer complexes from six species, including Homo sapiens, Mus musculus, E. coli, Bacteria, Eukaryota, and S. cerevisiae were kept in the datasets for training. The dimer datasets were then further screened by using sequence alignment. All the redundant proteins with over 50% sequence identity were removed from the dataset. In addition, the small proteins with lengths less than 50 amino acids were removed as well. The filtered protein dimer complexes were used as positive samples. Negative samples were generated by randomly pairing the chains. To maintain the balance of the training data, the sizes of the negative datasets were set to the same as the positive datasets. With the aforementioned filtering criteria and random pairing strategy, six single-species datasets and a mixed-species (including all six species) dataset were constructed (Table 1).

**Table 1.** Statistical data of multi-species dataset.

| Dataset | Positive | Negative | Total |
| --- | --- | --- | --- |
| Homo sapiens | 2690 | 2690 | 5380 |
| E. coli | 369 | 369 | 738 |
| Bacteria | 4975 | 4975 | 9950 |
| Eukaryota | 3420 | 3420 | 6840 |
| Mus musculus | 575 | 575 | 1150 |
| S. cerevisiae | 338 | 338 | 676 |
| Mixed-species | 10604 | 10604 | 21208 |

### Model implementation

The training was carried out by using PyTorch in Python. The RSPPI model was trained for 200 epochs using the Adam optimizer. The learning rate was set to 0.001 initially which exponentially decays by a gamma of 0.98 every epoch. The batch size was set to 1 during training. To avoid overfitting, the dropout technique was used, and the weight decay was set to 0.03 during training.

### Evaluation standard

The performance of the models was evaluated by classification accuracy, precision, sensitivity, F1-score (F1), and Matthews correlation coefficient (MCC). These parameters are defined below:

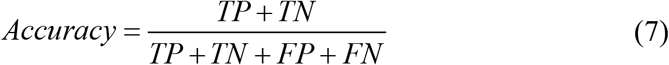

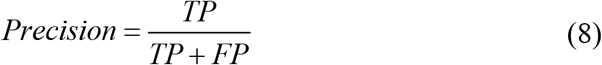

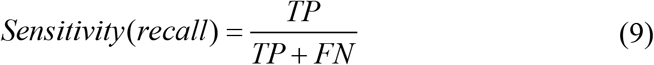

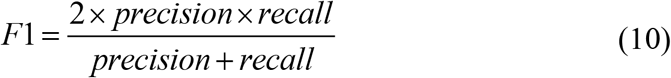

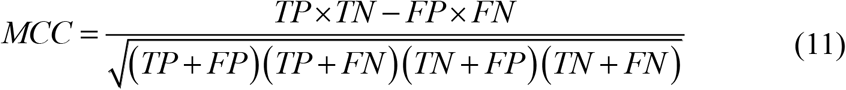

where TP, TN, FP, and FN stand for true positive, true negative, false positive, and false negative, respectively. A Receiver Operating Characteristic (ROC) curve indicates the performance of a classification model at all classification thresholds. The Area Under the ROC Curve (AUC) was also used to evaluate the model. In total, the above six evaluation specs were used to evaluate the RSPPI model.

## Results

From the ablation study, it was demonstrated the importance of the spatial structural information and SPP layer. Then the RSPPI was applied to the cross-species dataset and six single-species datasets. The trained model showed generally high precision both in the same species and cross species.

### Importance of the spatial structural information

To study the contribution of spatial structural and physicochemical information, we evaluated the performance of using three kinds of embedding features, i.e., using physicochemical features (hydrophobic and charge) only, using the structural feature only, and using all features. In the Mus musculus dataset, the performance of the model trained on the structural feature was better than that trained on the physicochemical features, while the model trained on all the features achieved the best performance (Fig. 6). Obviously, adding the structural feature will effectively improve the evaluation metrics, especially the precision of the model in the PPI prediction.

**Figure 6.**
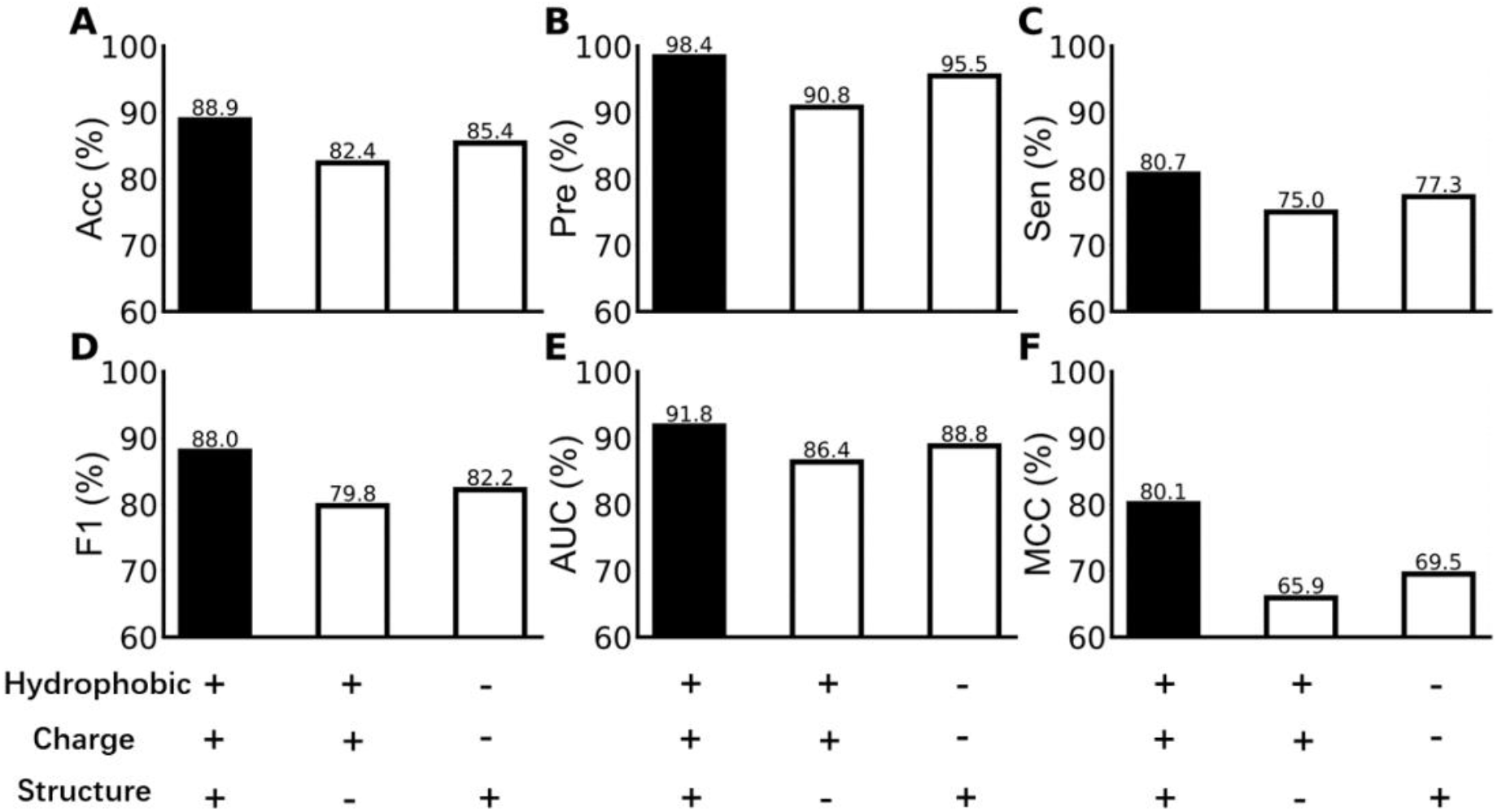
The performance of using three kinds of embedding features, including Accuracy (A), Precision (B), Sensitivity (C), F1-score (D), Area Under the ROC Curve (E), and Matthews correlation coefficient (F).

### Importance of Spatial Pyramid Pooling Layer

To avoid padding or cropping of protein, we equipped the networks with an SPP layer. To investigate the effect of the SPP layer, we compared our model with a fixed-length model trained on the E. coli and the Mus musculus datasets. In the fixed-length model, we used a similar model architecture, including six ResNet layers at the beginning and a fully connected layer at the end. In addition, all the proteins fed to the model were preprocessed to a fixed sequence length of 800 amino acids by padding the short proteins with zeros and cropping the long proteins. The batch size was set to 32 for the fixed-length training and other parameters were kept the same as the previous training with the SPP layer. Obviously, the model with the RSPPI achieved better performance than the fixed-length model in predicting the PPI on both the E. coli and Mus musculus datasets (Fig. 7).

**Figure 7.**
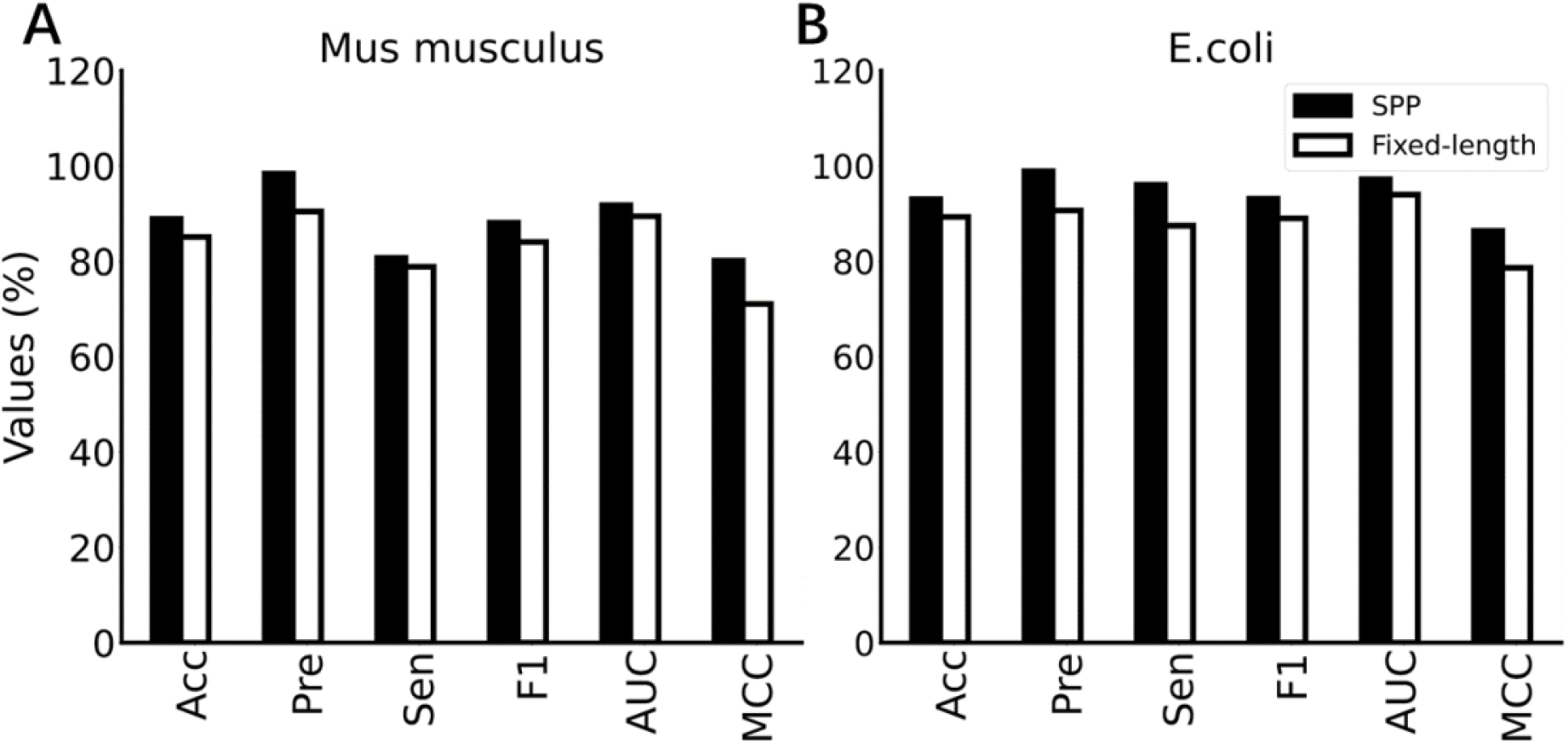
The performance comparison between models with and without the SPP layer. A. The performance comparison on the Mus musculus dataset. B. The performance comparison on the E.coli dataset.

### Evaluation of RSPPI on the intraspecies dataset

To test the performance of RSPPI on the intraspecies dataset, the 10-fold cross-validation was carried out on the Homo sapiens, Mus musculus, E. coli, S. cerevisiae, Bacteria, Eukaryota, and the mixed-species datasets, respectively. The mean evaluation parameters were calculated from the tenfold cross-validation (Fig. 8). In all types of species, the RSPPI exhibited high precisions, while showing acceptable performances in the accuracies, sensitivities, F1s, and AUCs. In all the single species datasets, RSPPI presented an average accuracy of ∼91%, a precision of ∼99%, a sensitivity of ∼87%, an F1 of ∼90%, an AUC of ∼93%, and an MCC of ∼82%. The model showed the best performance in the bacteria dataset with accuracy, sensitivity, and F1 all above 93%, an AUC above 97%, and a precision above 99%. In the Mixed species dataset, RSPPI presented an accuracy of 93%, a precision of 97%, an F1 of 92%, and an AUC of 92%. According to the evaluation parameters, the RSPPI model had good performance in all the tested species.

**Figure 8.**
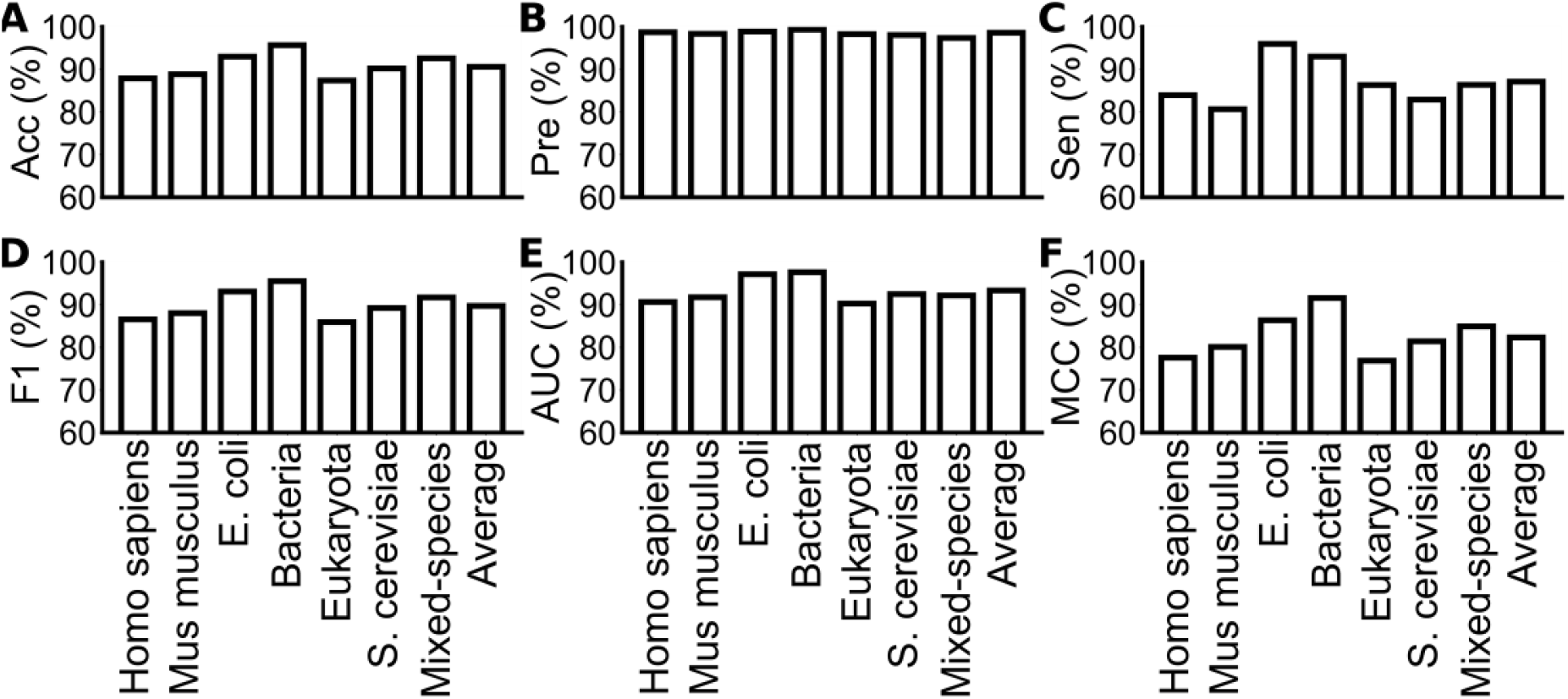
The performance of the RSPPI model evaluated on intraspecies datasets, including Accuracy (A), Precision (B), Sensitivity (C), F1-score (D), Area Under the ROC Curve (E), and Matthews correlation coefficient (F).

### The evaluation of the cross-species performance of RSPPI

To explore the generalization of RSPPI in predicting PPI, a cross-species validation was carried out, in which the model was trained with the Homo sapiens dataset and tested with all the other five species (Fig. 9). Surprisingly, our model presented a comparable performance in the cross-species validation comparing to the performance in the intraspecies evaluation, while most state-of-art PPI models failed to provide an acceptable prediction on cross-species dataset [16]. The Homo sapiens dataset trained model presented an average prediction accuracy of >90% on all the other five datasets, while the average precision of the intraspecies datasets is ∼98%. Among all the datasets, Eukaryota exhibited a relatively low accuracy of ∼87%, while the Bacteria dataset exhibited the highest accuracy of >95%. The Homo sapiens dataset trained RSPPI model learned the features of Homo sapiens PPIs. However, the model predicted the PPIs of the other species with similar performance as in the intraspecies evaluation, which suggested that the RSPPI combined the physicochemical properties and spatial structure information is able to predict the PPIs across species.

**Figure 9.**
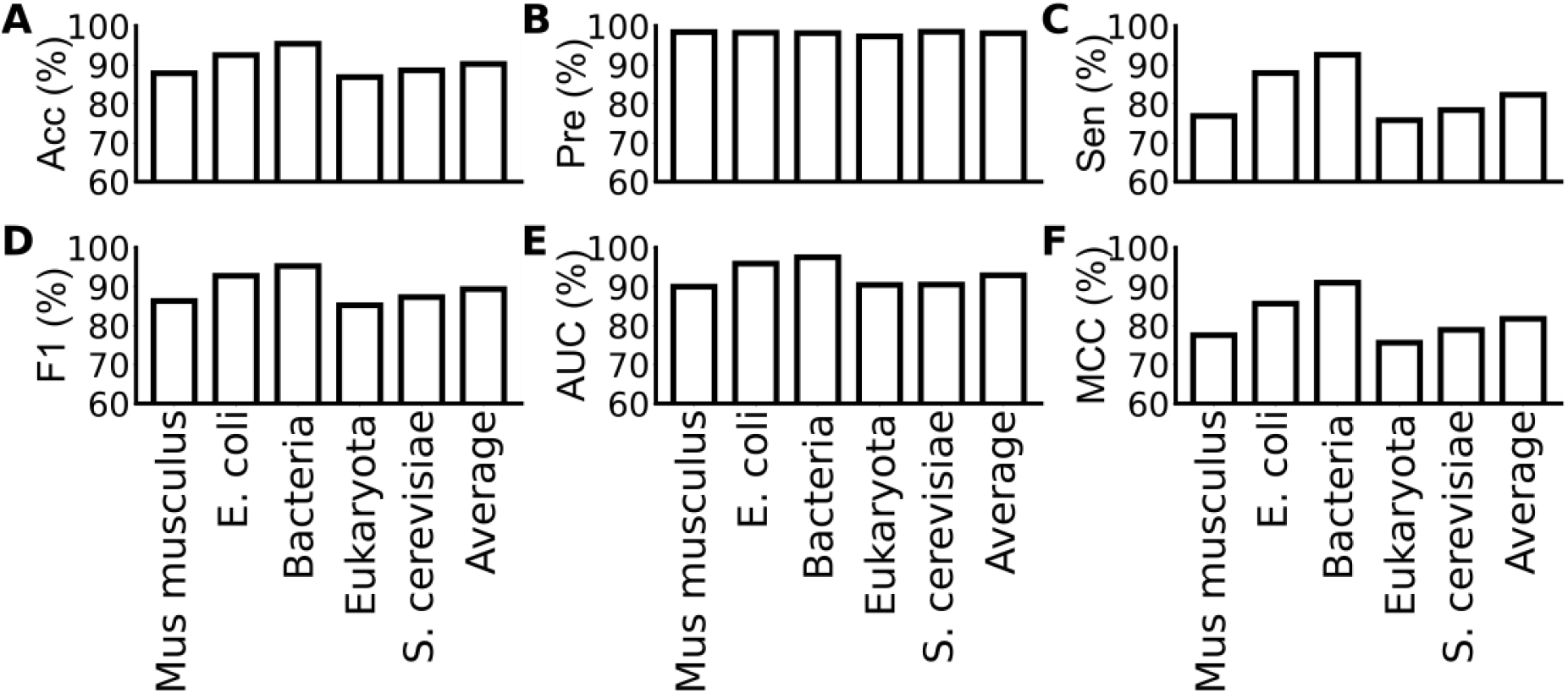
The performance of RSPPI evaluated with cross-species datasets, including Accuracy (A), Precision (B), Sensitivity (C), F1-score (D), Area Under the ROC Curve (E), and Matthews correlation coefficient (F).

### Comparison with the existing PPI prediction method

In order to compare our RSPPI model with other models, we listed the evaluation metrics of our model and five state-of-the-art methods, including KNN_LD [33], SVM_LD [11], PRED_PPI [34], PPI_MetaGO [35], Go2ppi_RF [20], trained with the PPIs information of the E. coli dataset side-by-side (Fig. 10). The KNN_LD, SVM_LD, and PRED_PPI models are based on the primary sequence of proteins, while the PPI_MetaGO and Go2ppi_RF models are based on the gene ontology (GO) information of proteins. AUC provides an aggregate measure of performance across all possible classification thresholds, in which a higher value represents the better classification ability of the model. MCC describes the correlation coefficient between the actual sample and the predicted sample, and its value close to 1 indicates that the prediction is very accurate. On the E. coli species dataset, RSPPI is superior to all these models in AUC and MCC. The RSPPI outperformed KNN_LD, SVM_LD, PPI_MetaGO, and GO2ppi_RF by 15.7%, 2.4%, 2.1%, and 2.0% in AUC, respectively, and outperformed PPI_MetaGO and GO2ppi_RF by 5.9% and 5.1% in MCC, respectively. In other metrics, RSPPI was one of the top 2 models, slightly behind SVM_LD in accuracy, sensitivity, and F1, and slightly behind KNN_LD in precision. Although SVM_LD ranked top 1 in three metrics, its performance in precision is the lowest among all models. Similarly, the top model, KNN_LD, concerning the precision, exhibited the worst performance in all the other metrics. Overall, our model performance is adequately good and well-balanced.

**Figure 10.**
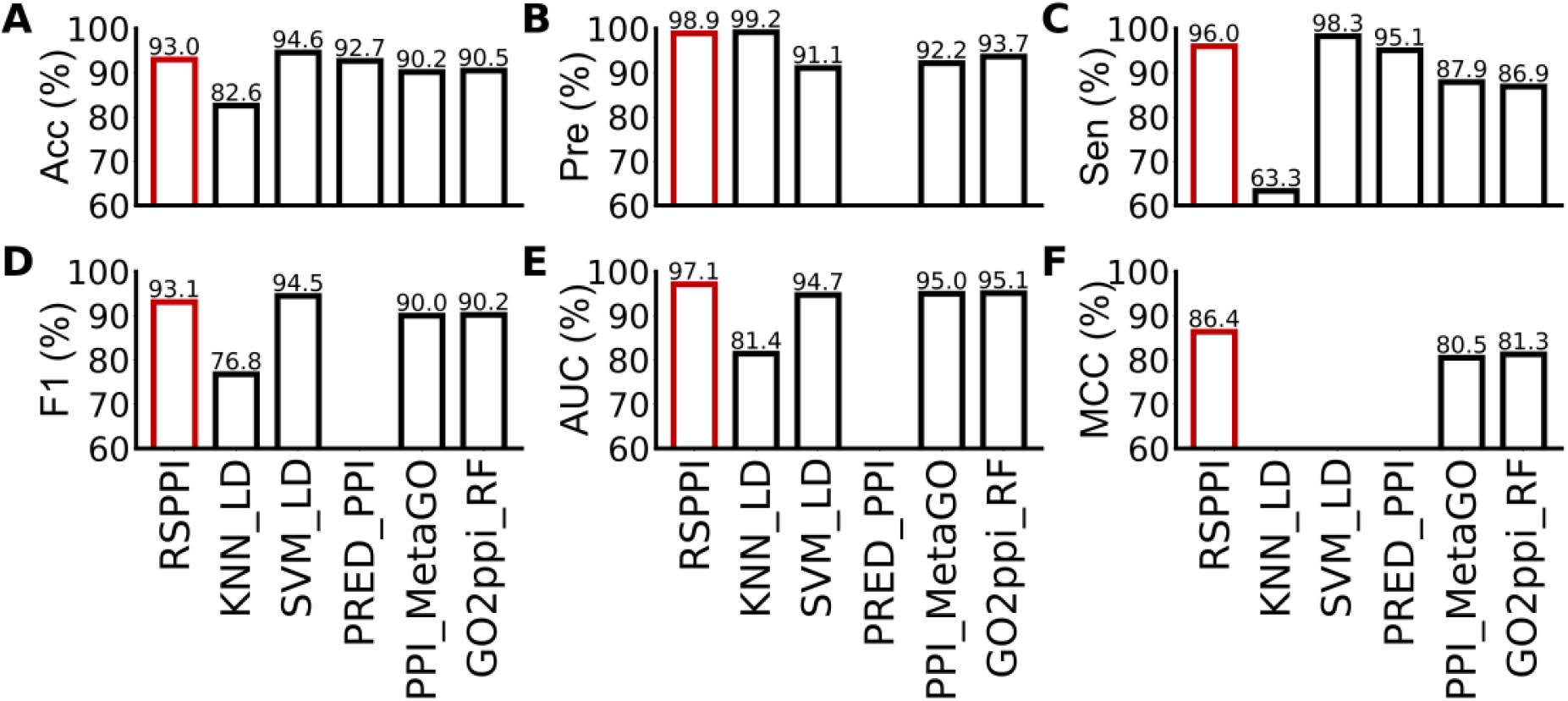
The performance comparison with other methods on the E. coli dataset, including Accuracy (A), Precision (B), Sensitivity (C), F1-score (D), Area Under the ROC Curve (E), and Matthews correlation coefficient (F).

Moreover, KNN_LD, SVM_LD, and three additional models, PCA_EELM [12], SVM_ACC [13], and SVM_PIRB [23] have been trained on S. cerevisiae. To further compare our model to these models, we summarized the evaluation metrics of our RSPPI model and the aforementioned five state-of-the-art models trained on S. cerevisiae side-by-side (Fig. 11). Among these models, KNN_LD, SVM_LD, PCA_EELM [12], and SVM_ACC [13] were based on the primary sequence of proteins, while SVM_PIRB [23] was based on the secondary structure. On the S. cerevisiae dataset, RSPPI led in both accuracy and precision and even showed great advantages in precision over all the other models, and was only left behind in sensitivity and AUC. The accuracy of RSPPI outperformed KNN_LD, SVM_LD, PCA_EELM, SVM_ACC, and SVM_PIRB by 4.0%, 1.6%, 3.2%, 0.9%, and 2.2%, respectively. By integrating physicochemical features and global spatial structure information, RSPPI achieved the best performance in most evaluation metrics.

**Figure 11.**
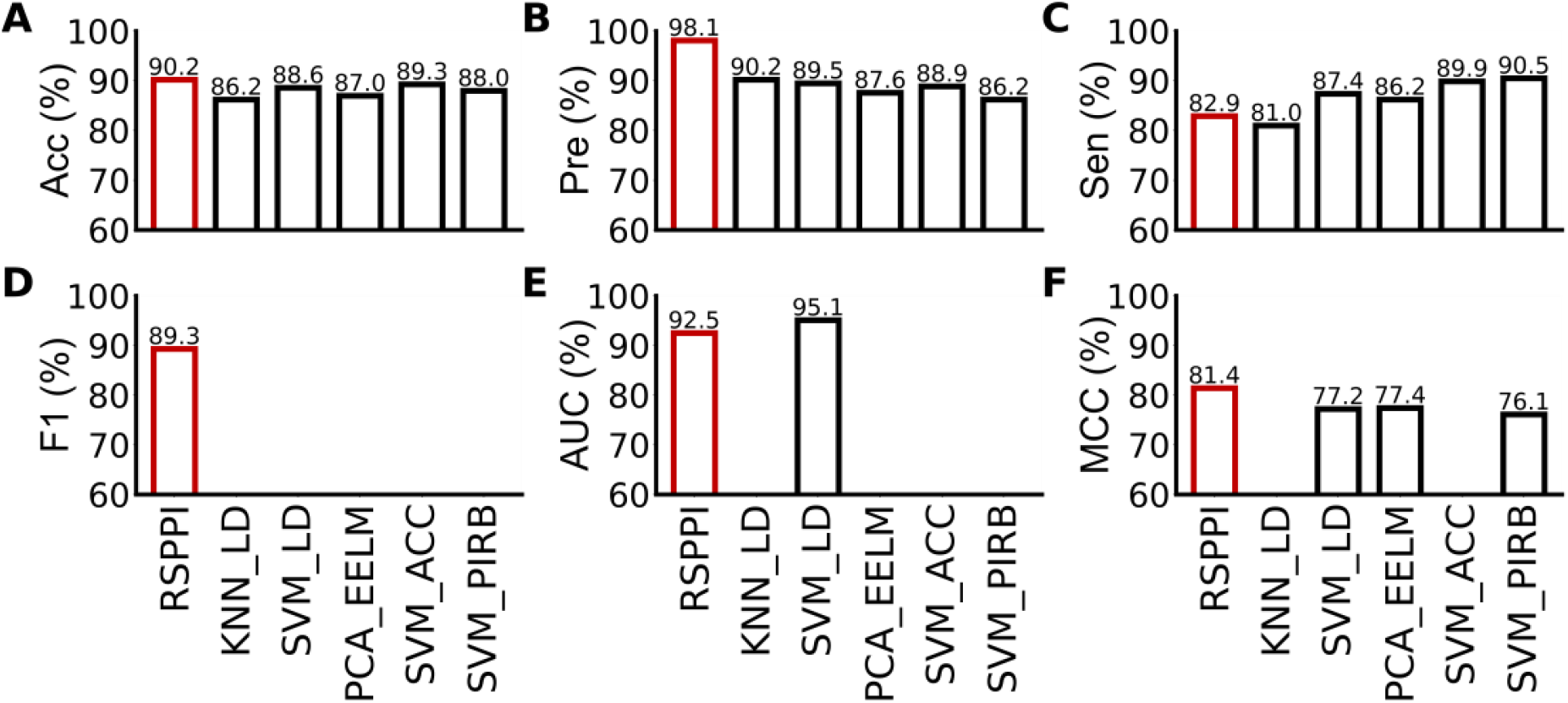
The performance comparison with other methods on the S. cerevisiae, including Accuracy (A), Precision (B), Sensitivity (C), F1-score (D), Area Under the ROC Curve (E), and Matthews correlation coefficient (F).

## Conclusion

In this paper, we propose RSPPI, a novel end-to-end deep learning method to predict PPIs. RSPPI learned from comprehensive feature profiles of proteins based on their physicochemical properties and spatial structure information. Systematic evaluations showed that RSPPI achieved good performance and high precision in both intraspecies and cross-species PPI prediction. The excellent performance on cross-species PPI prediction suggests the RSPPI has the potential to predict the less studied species based on current knowledge. As the structure and physicochemical compatibility are the basis of non-covalent PPIs, our RSPPI model based on spatial structure and physicochemical information showed better performance in most evaluation metrics, comparing to the sequence-based, GO-based, and secondary structure-based methods. In addition, by utilizing the Spatial Pyramid Pooling layer, RSPPI avoided the fixed-length requirement for the fully connected layer, thus reducing the artifact from padding or cropping. The RSPPI model provides a new strategy for PPI prediction by combining the spatial and physicochemical features of the protein.

## Availability of data and codes

The code, datasets, and trained model in this study are available in GitHub repository: https://github.com/Mechnobiology/RSPPI.

